# Misperceiving momentum: computational mechanisms of biased striatal reward prediction errors in bipolar disorder

**DOI:** 10.1101/2023.07.11.548610

**Authors:** Hestia Moningka, Wael El-Deredy, Richard P Bentall, Liam Mason

## Abstract

**Background:** Dysregulated reward processing and mood instability are core features of bipolar disorder that have largely been considered separately, and with contradictory findings. We sought to test a mechanistic account that proposes that, in bipolar disorder, there is an excessive tendency to enter recursive cycles in which reward perception is biased by signals that the environment might be changing for the better or worse.

**Methods:** Participants completed a probabilistic reward task with functional Magnetic Resonance Imaging. Using an influential computational model, we ascertained whether participants with bipolar disorder (*n* = 21) show greater striatal tracking of momentum-biased reward prediction errors (RPEs) than healthy controls (*n* = 21). We conducted psychophysiological interaction analyses to quantify the degree to which each group modulated functional connectivity between the ventral striatum and left anterior insula in response to fluctuations in momentum.

**Results:** In the bipolar disorder group, but not controls, the momentum-biased RPE model accounted for significant additional variance in ventral striatal activity beyond a standard mode of veridical RPEs. Compared to controls, the bipolar disorder group exhibited lower ventral striatal-left insular functional connectivity modulated by momentum-biased RPE. Moreover, this reduction in connectivity was more pronounced as a function of current manic symptoms.

**Conclusions:** Consistent with an existing theory, we found evidence that bipolar disorder is associated with a tendency for momentum to excessively bias striatal tracking of RPE signals. We identify impaired striatal-insular connectivity as a possible locus for this propensity. We argue that computational psychiatric approaches that examine momentary shifts in reward and mood dynamics have strong potential for new mechanistic insights and targets for intervention.

## Introduction

Mood instability is emerging as an important feature across psychiatric disorders including bipolar and depressive disorders, psychotic disorders, and personality disorder (Broome et al., 2015; Patel et al., 2015). It is characterised as a tendency to experience frequent oscillations of intense affect to a degree that it impacts one’s ability to regulate these changeable moods and their behavioural consequences (Marwaha et al., 2013). Mood instability predicts poorer clinical outcomes (Patel et al., 2015), including higher rates of relapse into bipolar mood episodes and rates of hospitalization (Gershon & Eidelman, 2015; Strejilevich et al., 2013). Mood instability persists even outside of overt mood episodes and predicts the development of future mood episodes (Broome et al., 2015). Despite growing recognition of its clinical importance, the mechanisms underlying mood instability in bipolar disorder are poorly understood.

A large body of neuroimaging work has examined the idea of dysregulated pursuit of reward in bipolar disorder; specifically the idea that patients are highly reactive to life events involving the attainment or failure to obtain reward (Nusslock & Alloy, 2017; Urošević et al., 2008). Task-based functional Magnetic Resonance Imaging (fMRI) studies have revealed differences in reward sensitivity in bipolar disorder; however, the direction of these effects is highly inconsistent. Even in studies of patients outside of major mood episodes, comparable numbers of studies find decreased reward sensitivity (Johnson et al., 2019; Schreiter et al., 2016; Yip et al., 2015) as increased (Mason et al., 2014; Nusslock et al., 2012). One issue is that studies typically rely on averaging responses to reward and punishments over entire experiments, whereas the clinical literature suggests that the relationship between mood and reward may be more dynamic. Another issue is that reward-based decision-making is a multi-process and recursive phenomenon, comprising anticipatory processes (including expectation), outcome evaluation and learning and other signals that carry over to the next decision (Dreher, 2013; O’Doherty et al., 2017). Crucially, whilst existing studies recognize that these reward processes impact mood, there is emerging evidence demonstrating that changes in mood in turn also impact on reward and decision-making processes (Eldar & Niv, 2015; Vinckier et al., 2018).

Computational approaches with a focus on generative models that link unobservable brain states to experimental measurements are well-suited to derive mechanistic explanations of neuroimaging data (Stephan et al., 2015) and could therefore capture more precisely the dynamics of mood instability in bipolar disorder. A recent neurocomputational account of bipolar disorder proposed that strong and changeable moods may be underpinned by a propensity for biased perception of reward value, resulting in recursive cycles in which mood, expectations and behaviour oscillate between extremes (Mason et al., 2017). This explanatory account builds on recent computational fMRI work, which found that violations of expectations in relation to outcomes (i.e. reward prediction errors, RPEs) drive transient mood fluctuations (Eldar & Niv, 2015; Vinckier et al., 2018).

Eldar and Niv (2015) showed that healthy individuals with higher trait mood instability exhibited increases in striatal activation to reward outcomes when in a positive mood state and decreases when in a negative mood state. Across two experiments, they demonstrated support for a generative model in line with the framework put forward by Eldar and colleagues (2016). This computational account therefore holds great promise in capturing the polar extremes of mood fluctuations such as those seen in bipolar disorder. Under this model, momentary mood is formalised as an integration of recent RPEs, representing an overall momentum of reward, in which a higher momentum of rewards and losses engenders a positive and negative mood state respectively. This quantity was shown to bias the perception of subsequent reward outcomes, and better accounted for striatal activation than the standard (unbiased) reinforcement learning model. This model was further corroborated in a subsequent study probing the mechanisms of action of antidepressant medication (Michely et al., 2020). In related work, Vinckier and colleagues (2018) also demonstrated that momentary mood biases the valuation of potential rewards and losses during decision-making. Their work also implicated anterior insula in representing trial-by-trial variation in recent RPE history (“mood fluctuations”), dovetailing with earlier work in which Rutledge and colleagues (2014) localised self-reported mood to anterior insula activation. Collectively, there is compelling evidence that mood fluctuations can modulate RPE signals represented in the ventral striatum. These models have great potential for better understanding and targeting the mechanisms underlying mood instability, but have not yet been utilised with clinical populations (Mason et al., 2017).

The present study puts to the test the central hypothesis we have outlined previously (Mason et al., 2017); specifically, individuals with bipolar disorder have an elevated propensity to enter recursive cycles of biased striatal RPEs. We expected that, as in Eldar and Niv (2015), both groups would show tracking of a signal representing the recent history of outcomes (what we term here as “momentum”) and that this signal would be reflected in the anterior insula (Rutledge et al., 2014; Vinckier et al., 2018). We hypothesised, however, that only the bipolar disorder group would there be a recursive cycle in which momentum biases the perception of subsequent rewards. Specifically, we predicted that striatal activations in the bipolar disorder group, but not the healthy controls, would track these “momentum-biased” RPEs. Finally, we sought to isolate the mechanism by which momentum biases reward perception. We predicted that this propensity would be underpinned by stronger coupling between the ventral striatum and anterior insula in the bipolar disorder group compared to healthy controls. We predicted that the differences in ventral striatal activation and connectivity would be larger in participants with higher levels of residual mood symptoms.

## Materials and Methods

### Participants and power calculation

Participants were people with bipolar disorder out of a major mood episode (*n* = 21) and healthy controls (*n* = 21) matched by age, sex and level of education (Table 1). The clinical group was recruited from local mental health trusts and specialist affective disorder clinics in Greater Manchester, United Kingdom, and the control group were recruited from the general community, as described in a previous study (Mason et al., 2014).

**Table 1.** Participant demographics and behavioural data. *Data from 2 participants are missing and not included. BD-I = Bipolar I Disorder; BD-II = Bipolar II Disorder; GAD = Generalised Anxiety Disorder; AUD = Alcohol Use Disorder; SUD = Substance Use Disorder; OCD = Obsessive-Compulsive Disorder; HAMD-17 = Hamilton Depression Rating Scale; MAS = Bech-Rafaelsen Mania Scale; BIS-11 = Barratt Impulsiveness Scale; SSRI = Selective Serotonin Reuptake Inhibitor; SNRI = Serotonin-Norepinephrine Reuptake Inhibitor.

|  | Bipolar group<br>(n = 21) | Healthy control group<br>(n = 21) | Statistic | p-value |
| --- | --- | --- | --- | --- |
| Age, M (SD)* | 35.95 (8.34) | 33.25 (9.32) | $t(38) = -.97$ | .34 |
| Female, n (%) | 11 (52.4%) | 9 (42.9%) | $\chi^2(1) = .38$ | .76 |
| Years of education, M (SD)* | 14.08 (2.47) | 14.70 (2.29) | $t(38) = .83$ | .41 |
| Primary diagnosis, n |  |  |  |  |
| BD-I | 18 |  |  |  |
| BD-II | 3 |  |  |  |
| Current comorbidity, n* |  |  |  |  |
| GAD | 2 |  |  |  |
| Lifetime diagnoses, n* |  |  |  |  |
| AUD/SUD | 10 |  |  |  |
| Panic disorder | 4 | 1 |  |  |
| GAD | 2 |  |  |  |
| OCD | 1 |  |  |  |
| Medications, n* |  |  |  |  |
| Lithium | 8 |  |  |  |
| Valproate | 5 |  |  |  |
| Lamotrigine | 2 |  |  |  |
| SSRI | 3 |  |  |  |
| SNRI | 3 |  |  |  |
| Benzodiazepine | 1 |  |  |  |
| z hypnotic | 3 |  |  |  |
| None | 4 |  |  |  |
| HAMD-17, M (SD) | 3.83 (2.96) | .60 (1.04) | $t(24.90) = -4.73$ | $p < .001$ |
| MAS, M (SD) | 3.55 (3.09) | .38 (1.11) | $t(25.05) = -4.43$ | $p < .001$ |
| BIS-11, M (SD) | 62.62 (6.00) | 76.95 (9.94) | $t(32.85) = -5.66$ | $p < .001$ |

The main inclusion criteria were 18 – 45 years of age, a weekly alcohol intake of below 26 units and no substance use four months before the study. The Structured Clinical Interview for DSM-IV Axis I Disorders (SCID; First et al., 2005) was used to confirm the diagnosis of bipolar disorder and screen healthy control participants. Individuals who did not meet threshold for either manic or depressive episodes (i.e. out-of-episode) for two months before the study were included. Individuals were excluded if they had received antipsychotic medication six months before the study. Residual manic and depressive symptoms were assessed using the 12-item Bech-Rafaelsen Mania Scale (MAS; Bech, 1995) and 17-item Hamilton Depression Rating Scale (HAMD-17; Hamilton, 1960) respectively. Trait impulsivity was assessed using the 11-item Barratt Impulsiveness Scale (BIS-11; Patton et al., 1995). All participants provided written informed consent.

Power analysis was informed by Eldar and Niv (2015)’s study, which compared striatal and ventromedial prefrontal cortex activity between healthy participants exhibiting low versus high trait mood instability, as defined by a median split on the Hypomanic Personality Scale (Eckblad & Chapman, 1986). Using a similar reward-based task, they found a significant difference in striatal activations between participants with high and low mood instability with an effect size of *d* = .90, which we used to determine the implied power in our study. We conducted a power calculation using G*power (Faul et al., 2007) and yielded 89% implied power with our sample size of 42 participants at an alpha level of .05. Given that our study examines striatal activation between individuals with bipolar disorder and healthy controls, the effect size obtained should be significantly larger than those obtained in Eldar and Niv (2015)’s sample of healthy participants.

### Task

Participants completed a variant of a validated Roulette task (van Eimeren et al., 2009), in which reward probability and magnitude were independently manipulated. Each trial consists of three stages: choice, anticipation and outcome (Figure 1). During choice, participants chose between four options that confer the same probability of reward. In low probability trials (25% chance of reward), participants selected one of four individual colours that made up the Roulette wheel. In high probability trials (75% chance of reward), they selected between four sets of three colours and won if the Roulette wheel stopped on any one of the three colours. The stake on offer (reward magnitude) was also presented prior to choice, with equal numbers of low (£3) and high (£9) magnitude trials. During anticipation, the Roulette wheel spun (3-4 seconds). At outcome, the wheel stopped spinning with the location of the Roulette ball indicating whether the participant had won or lost the amount of money at stake.

**Figure 1.**
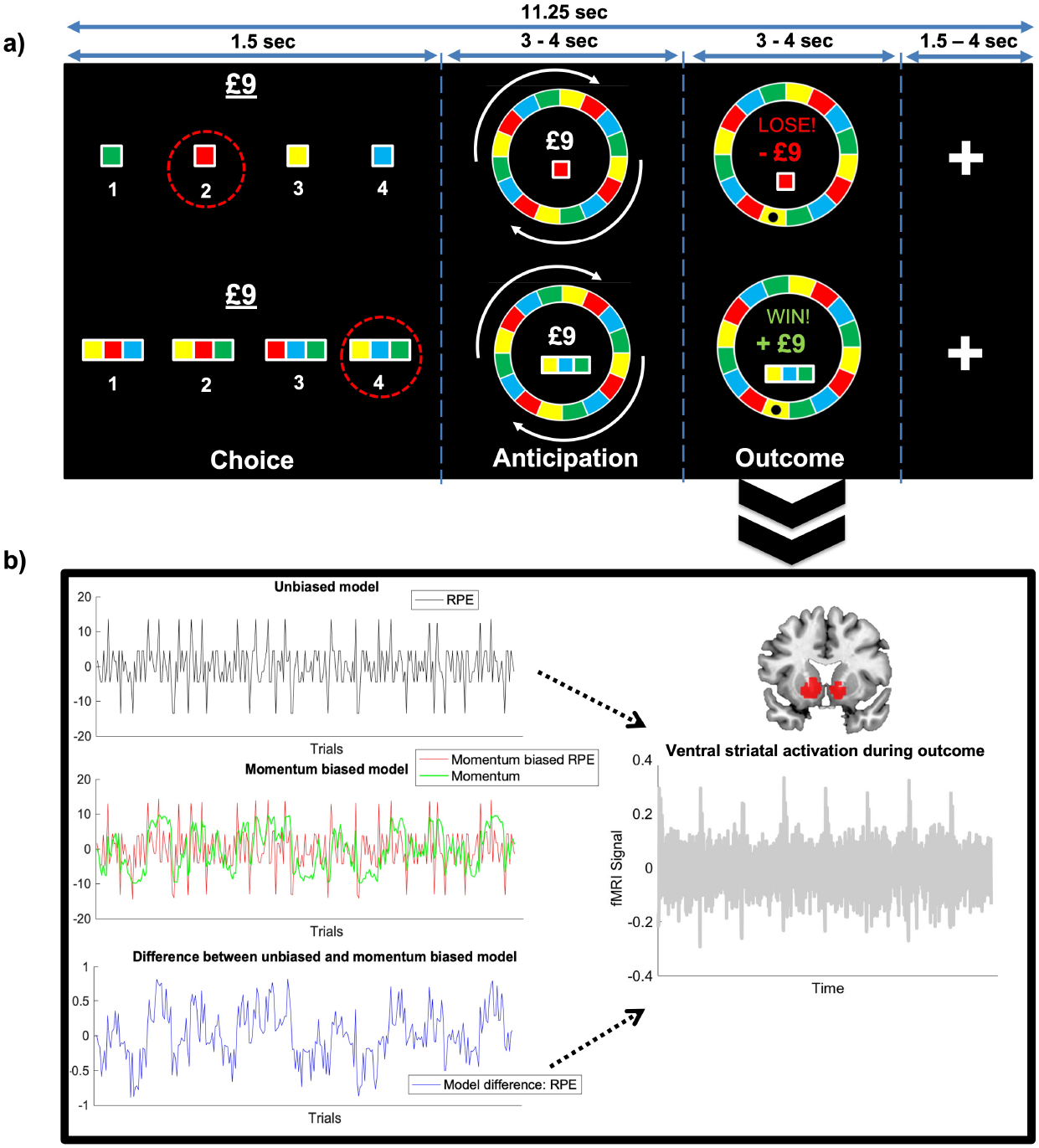
Schematic of the Roulette Task and Model-based fMRI. a) Roulette Task. Participants placed bets on which colour would win in a Roulette spin, alternating between trials in which £3 or £9 was at stake. The red dotted circles during ‘Choice’ correspond to an example choice a participant could make under a low (25%) or high (75%) probability of reward. After betting, the wheel began spinning (anticipation) before stopping and participants received feedback (outcome). b) Model-based fMRI with parametric modulators for trial-wise reward prediction error (RPE) during outcome evaluation. The RPE time series was generated under a standard, unbiased model, and under a recursive biased model in which the momentum of recent outcomes impacted the perception of subsequent outcomes. Adapted from Mason et al. (2014).

Participants were instructed to respond within the fixed duration of the choice phase; otherwise, a random choice would be automatically selected for that trial. They were informed that they would be paid the actual winnings following task completion. Participants completed 272 trials across eight blocks, each lasting approximately six minutes.

### FMRI acquisition and preprocessing

Standard fMRI data acquisition of functional images was conducted as previously reported in Mason et al. (2014). Echo-planar image sequence with repetition time = 2450ms, echo time = 25ms, flip angle = 90°, slices = 30 in ascending order, slice thickness = 4mm, in-plane resolution = 1.5 × 1.5mm and a standard field-of-view was acquired from a 1.5T Phillips scanner. Data were processed using SPM12 (Wellcome Centre for Human Neuroimaging, University College London) and Matlab R2019b. Each participant’s functional images were motion-corrected using a six-parameter rigid-body transformation to the mean image and slice-time-corrected to the middle slice. Functional images were co-registered with each participant’s structural image, spatially normalised to the Montreal Neurological Institute (MNI) standard template and smoothed using an 8mm Gaussian kernel. Intrinsic autocorrelations were accounted for by AR(1) and low frequency drifts were removed via the 128s high pass filter. The ArtRepair toolbox (Mazaika et al., 2009) was used to minimise the impact of artifacts via the interpolation of outlier volumes, which resulted in the inclusion of fMRI data from all 42 participants.

### Model-based functional MRI

First-level analyses were a general linear model (GLM), implemented in SPM12. FMRI blood-oxygen-level-dependent (BOLD) responses in each run for each participant were modelled using regressors representing the three task conditions (Choice, Anticipation and Outcome), two parametric modulators modelled at Outcome (trial-wise unbiased RPE and recursively biased RPE) and six motion-realignment parameters to reduce residual effects of motion, resulting in a total of 11 regressors.

We generated trial-wise unbiased and momentum-biased RPE values generated from Eldar and Niv (2015)’s computational model, where momentary mood is formalised as the accumulation of recent outcomes (i.e. momentum of RPE) and influences the trial-wise RPE. Specifically, we computed unbiased RPEs from the expected values and outcomes received by participants. We then generated momentum-biased RPEs under a generative model in which recent RPEs are allowed to summate, and bias the perception of subsequent outcomes (Figure 2). Given evidence of impaired reinforcement learning in bipolar disorder (Bart et al., 2021; Urošević et al., 2018), we utilised a task that removed learning, precluding fitting exact individual participant parameters. Instead, we sought to quantify the extent that ventral striatal activations were consistent with a ‘momentum-biased’ agent, utilising the group average momentum bias parameter in participants with elevated hypomanic personality traits, previously reported by Eldar and Niv (2015) (see Supplementary Materials). This approach accords with recommendations for model-based fMRI (Wilson & Niv, 2015), which showed that effects when regressing RPE time series generated under a single average parameter for all participants are comparable to time series generated under individual parameter estimates.

**Figure 2.**
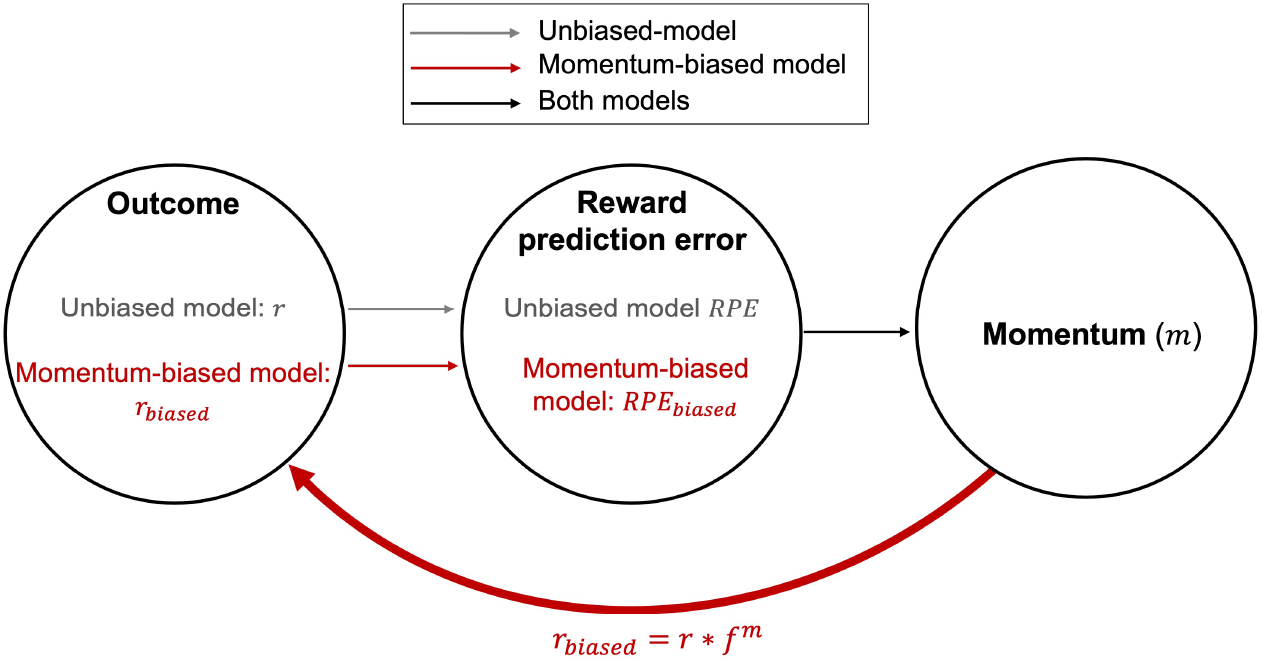
Illustration of unbiased and momentum-biased models.

Following Eldar and Niv (2015)’s approach, momentum-biased RPE was calculated as the difference between unbiased and momentum-biased estimates, to avoid regressor collinearity. Regressors were convolved with a canonical haemodynamic response function. Individual contrast images of interest (per-trial estimates of unbiased and momentum-biased RPE) were computed and passed to a second-level GLM to examine within-group and between-group activations. We additionally included model-estimated values of momentum as a parametric modulator in a separate GLM to corroborate that this signal is represented in the anterior insula, consistent with previous studies localising momentary mood to anterior insula activations (Rutledge et al., 2014; Vinckier et al., 2018).

### Striatal tracking of momentum-biased RPE

Analyses utilised a region of interest (ROI) approach. To define the ROI for ventral striatum, we followed the approach of Eldar and Niv (2015) and included all grey-matter voxels within the bilateral ventral striatum that responded to ‘reward’ outcomes versus ‘loss’ outcomes contrast (at a family-wise-error (FWE) corrected threshold of *p*FWE < .05). We also extracted activations from an insula ROI, defined as 8mm spheres around the peak coordinates for the bilateral anterior insula reported by Vinckier and colleagues (2018) (left: x = - 30, y = 22, z = -6; right: x = 32, y = 20, z = -6). Parameter estimates representing the mean activity across all voxels in these ROIs for each participant were extracted for each contrast of interest using MarsBar (http://marsbar.sourceforge.net/) using standard threshold of *p* < .05. For the ventral striatum ROI, the contrast of interest was the parametric modulation of outcome-locked activation by trial-by-trial momentum-biased RPE values (i.e. the difference between unbiased and momentum-biased estimates). For the anterior insula ROIs, the contrast was the parametric modulation by trial-by-trial momentum, applying Bonferroni correction for these two tests.

To test for associations with residual mood symptoms, manic and depressive symptoms were entered as predictors into a regression analysis in bipolar participants. Trait impulsivity was accounted for in this analysis, given that this had previously been shown to impact reward activations in this dataset (Mason et al., 2014). Log transformation was used for non-normal variables.

### Functional connectivity of striatal momentum-biased RPE activations

To examine ventral striatal-insular connectivity as a locus of momentum biasing reward perception, we quantified the extent that outcome-locked functional connectivity between these two regions was modulated by momentum-biased RPEs. We used generalised psychophysiological interaction analysis (gPPI; http://www.nitrc.org/projects/gppi) (McLaren et al., 2012) with seed region in our ventral striatum ROI. At first-level, the fMRI timeseries for outcome-locked activations was extracted from the seed region for each task run then multiplied by the task regressor of interest to form the PPI interaction terms. Second-level analyses were conducted with the gPPI images to identify within- and between-group connectivity between the ventral striatum and anterior insula using MarsBar, as described above. The contrast of interest for the functional connectivity analyses was the parametric modulation of outcome-locked activation by trial-by-trial momentum-biased RPE values (i.e. the difference between unbiased and momentum-biased estimates).

## Results

Participant demographics and behavioural results are reported in Table 1. Compared to healthy controls, participants with bipolar disorder showed higher levels of manic (Bech-Rafaelsen Scale; MAS) and depressive symptoms (Hamilton Depression Rating Scale; HAMD-17) and trait impulsivity (Barratt Impulsiveness Scale; BIS-11).

### Striatal tracking of momentum-biased RPE

Ventral striatal activation during the anticipation and outcome stage is reported elsewhere (Mason et al., 2014) and is both greater in participants with bipolar disorder than healthy controls.

Consistent with prior literature, bilateral ventral striatal activation tracked unbiased RPE across bipolar disorder [*t*(20) = 6.16, *p* < .001] and control [*t*(20) = 6.04, *p* < .001] groups.

Bilateral ventral striatal activation tracked momentum-biased RPE only in bipolar participants [*t*(20) = 2.61, *p* = .0063] and not healthy controls [*t*(20) = .78, *p* = .22] (Figure 3), partially corroborating our hypothesis. The bipolar disorder group additionally trended towards showing greater tracking of momentum-biased RPE than healthy controls, but this did not reach significance [*t*(40) = 1.46, *p* = .075].

**Figure 3.**
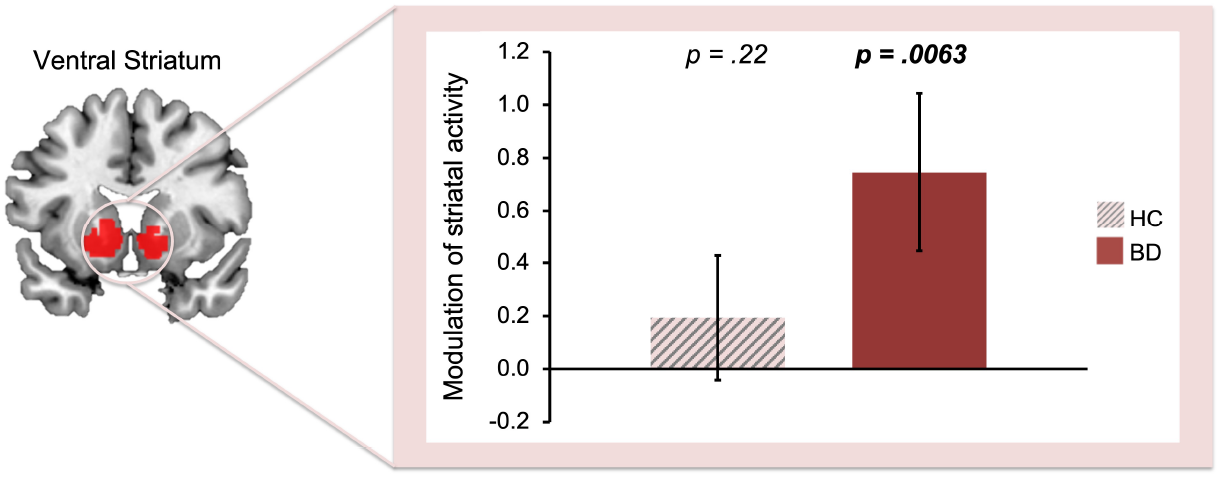
Modulation of outcome activity by momentum-biased RPE in the ventral striatum. Only the bipolar disorder group tracked momentum-biased RPE (error bars: standard error of the mean). HC = Healthy control group; BD = Bipolar disorder group.

This momentum-biased RPE modulated ventral striatal activity was not predicted by levels of manic and depressive symptoms in the bipolar disorder group (Supplementary Materials, Table S1).

Across both groups, we confirmed that momentum (i.e. the aggregate of recent outcomes; see Materials and Methods) was tracked in the anterior insula [left anterior insula selectively: *t*(41) = 2.11, *p* = .021, not right anterior insula: *p* = .31; Bonferroni-corrected *p*-value threshold is *p* = .025] (Figure 4). As expected, we did not find a group difference in the degree to which anterior insula tracked momentum (*p* ≥ .41).

**Figure 4.**
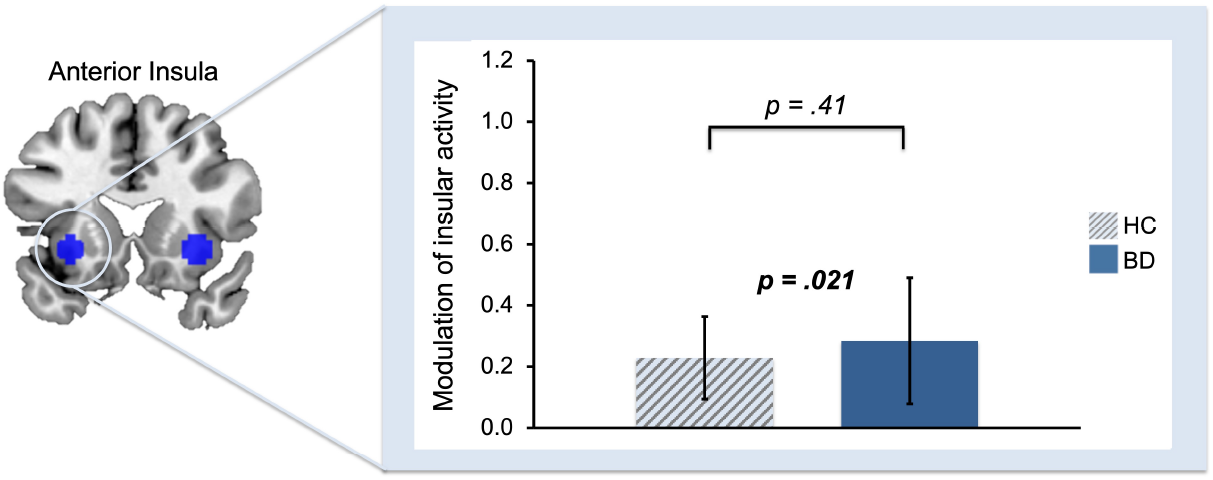
Activation in left anterior insula tracks the degree of momentum of RPEs. Across both groups, momentum was tracked by the left anterior insula (*N* = 42, *p* = .021) and this was not significant between groups (*p* = .41). Error bars reflect standard error of the mean. HC = Healthy control group; BD = Bipolar disorder group.

### Modulation of striatal connectivity by momentum-biased RPE

Next we sought to understand why ventral striatal activations tracked momentum-biased RPEs only in the bipolar disorder group (Figure 3) and whether this is accompanied by differences in the integration of the momentum signals we identified in left anterior insula activation. To do this, we quantified the degree to which the coupling between the ventral striatum and anterior insula is modulated by fluctuations in momentum. We focused on the left anterior insula, given that there was left-lateralised selectivity in tracking momentum.

We found that the bipolar disorder group diverged from healthy controls in how their ventral striatal connectivity with the left anterior insula was modulated by momentum-biased RPE [*t*(40) = 3.38, *p* = .00047]. This group difference, however, was in the opposite direction to our hypothesis. Whereas healthy controls showed a significantly stronger positive modulation of ventral striatal connectivity with the left anterior insula [*t*(20) = 3.24, *p* = .0012], bipolar participants showed modulation in the opposite direction [*t*(20) = 1.78, *p* = .041]. Moreover, the divergent negative coupling was greater in those with elevated residual manic symptoms (Figure 5) and as a function of trait impulsivity [MAS: *β* = -.45, *t* = -2.08, *p* = .05; BIS-11: *β* = .47, *t* = 2.26, *p* = .04] (Supplementary Materials, Table S2).

**Figure 5.**
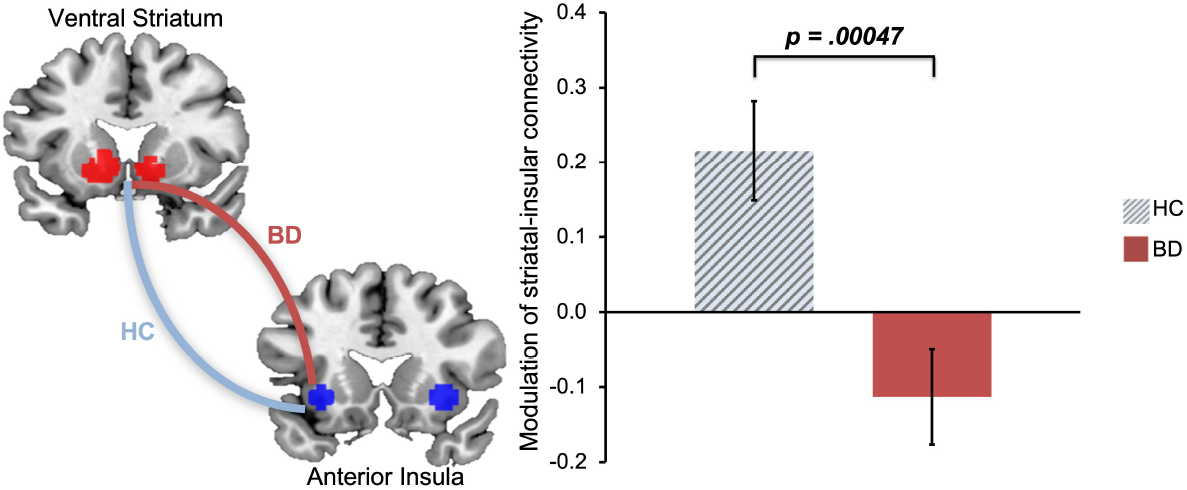
Divergent modulation of ventral striatal-left anterior insular functional connectivity by momentum-biased RPE in bipolar disorder group compared to healthy controls. Functional connectivity between bilateral ventral striatum and left anterior insula during outcome (error bars: standard error of the mean). The blue line represents modulation in the healthy control group (HC) and the red line represents modulation in the bipolar disorder group (BD).

For specificity, we confirmed that ventral striatal-left anterior insular connectivity was not modulated by unbiased RPE in either group (*p* ≥ .13).

## Discussion

This study put to the test the theory that bipolar disorder is underpinned by a propensity for reward perception to be biased by fluctuations in momentum of recent reward prediction errors (RPEs).

Consistent with the hypothesis put forward in our prior work (Mason et al., 2017), we found evidence that ventral striatal RPEs are excessively modulated by momentum in individuals with bipolar disorder (Figure 3). Moreover, bipolar participants showed divergent modulation of ventral striatal-left anterior insular connectivity, the extent of which increased as a function of current manic symptoms.

Consistent with prior work, we found that both bipolar disorder patients and healthy controls tracked the momentum of recent reward prediction errors (Figure 4). This quantity, previously shown to capture momentary mood (Eldar & Niv, 2015), was tracked by anterior insula, a region implicated in two other studies examining the intersection of mood and reward processes (Rutledge et al., 2014; Vinckier et al., 2018). However, only the bipolar disorder group showed evidence that this momentum quantity additionally biased RPE signals in the ventral striatum (Figure 3). Thus, we found evidence for the hypothesis that a bidirectional relationship between mood and reward processes, exists in bipolar disorder, such that fluctuations in momentum bias the perception of subsequent outcomes in a recursive cycle. We have previously argued, through simulations, that this mechanism is sufficient to drive the oscillatory changes in mood that culminate in manic and depressive episodes (Mason et al., 2017).

We sought to deepen this insight by asking how momentum might impact striatal RPEs in bipolar disorder. We reasoned that this would come about through excessive coupling between the anterior insula and ventral striatum, consistent with an over-propagation of the momentum signal. We found that striatal-insular functional connectivity diverged in bipolar disorder, but this was due to stronger negative coupling between these regions as a function of momentum-biased reward prediction errors. Thus, our findings suggest that in periods of upward momentum (better-than-expected outcomes), striatal-insular coupling becomes stronger in healthy controls and weaker in participants with bipolar disorder.

Whilst unexpected, one interpretation of the connectivity results is that greater coupling with the anterior insula serves to contextualise (rather than bias) reward perception. By holding a running total of momentum alongside new outcomes, this could suppress signals that would otherwise provide recursively confirmatory evidence that the environment is changing. In bipolar patients, the failure to contextualise, through strengthened positive connectivity during periods of upward momentum (better-than-expected outcomes), could increase susceptibility to misperceive upward momentum and to act on it, further escalating mood. Consistent with this possibility, we found that this divergent connectivity to be even stronger in patients with higher levels of manic symptoms (Figure 6), consistent with over-confidence and risk-taking during (hypo)mania (Ramírez-Martín et al., 2020; Savitz et al., 2008). This interpretation remains speculative until corroborated with further work, although we did confirm that the striatal-insular coupling was specifically modulated by momentum-biased RPEs and not unbiased RPEs.

**Figure 6.**
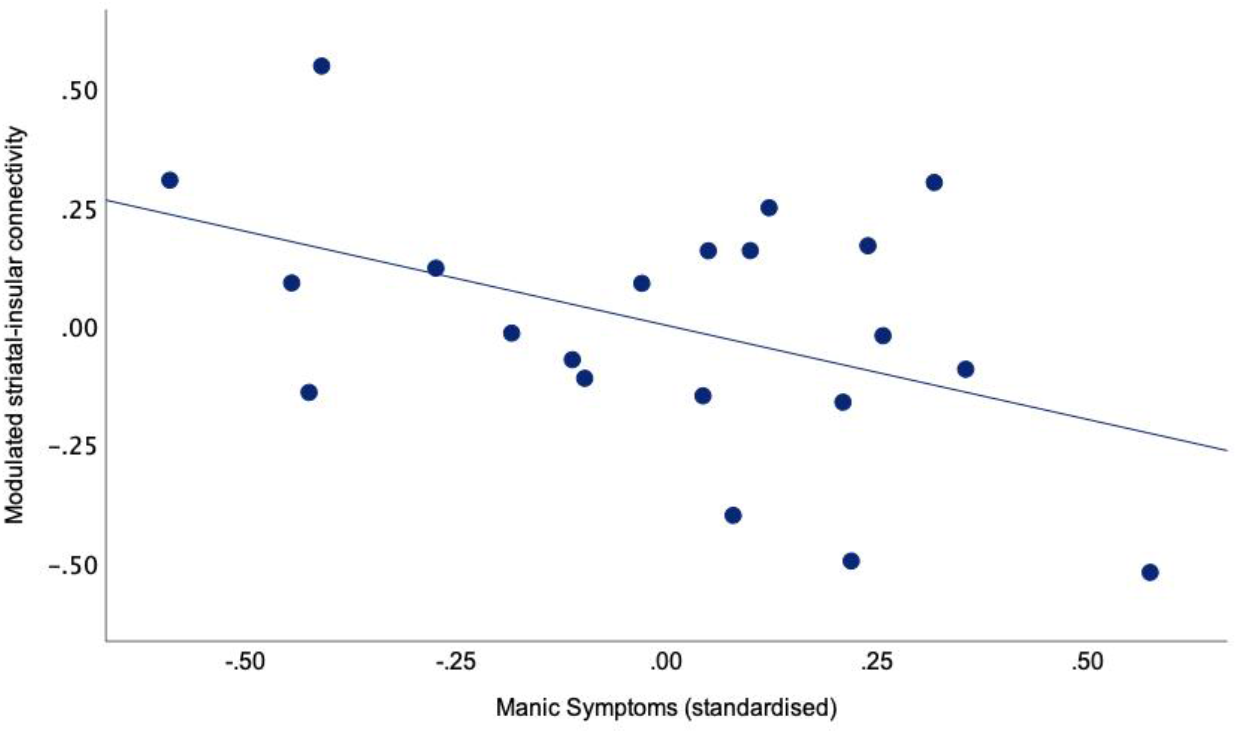
Stronger inverse modulation of ventral striatal-left anterior insular connectivity as a function of residual manic symptoms in the bipolar disorder group. Partial association from regression model of functional connectivity between bilateral ventral striatum and left anterior insula modulated by momentum-biased RPE. Log transformed scores from Bech-Rafaelsen Mania Scale.

Intriguingly, we observed that higher levels of trait impulsivity (BIS-11) offset the inverse striatal-insular coupling we observed in the bipolar disorder group. Within the above interpretation, trait impulsivity offsets the tendency to integrate momentum of past rewards when evaluating current outcomes. Impulsivity has long been conceptualised as a myopic focus on immediate rewards or a failure to integrate past temporal horizon (MacKillop et al., 2016; Ross et al., 2015) including disproportionate striatal activations for decisions involving immediate rewards (McClure et al., 2004). Previous analyses using the current dataset found impulsivity to predict reduced reactivity to safe (high-probability) rewards beneficial for longer-term goals (Mason et al., 2014). In addition, the poor sustained attention, non-planning and sensation-seeking facets of impulsivity tapped by BIS-11 may be less relevant to momentum-biased reward processing. Other measures that tap emotion-related impulsivity (i.e. engaging in rash behaviour following positive or negative emotions) have been highlighted as more relevant to mood instability in bipolar disorder (Bøen et al., 2015; Muhtadie et al., 2014). Consistent with this notion, we argue that we could be picking up a distinct dimension of impulsivity (beyond inattentive or non-planning impulsivity) in bipolar disorder, which might be an epiphenomenon of being excessively prone to momentum.

A limitation of our study is that whilst we assessed current mood symptoms on the day of the scan, we did not have ratings of momentary mood during the task. This would be useful to confirm that momentum corresponds to momentary mood, as found previously in participants with elevated hypomanic personality traits (Eldar & Niv, 2015) and in general population samples (Vinckier et al., 2018). Consistent with previous reports that mood instability perseveres outside of mood episodes (Broome et al., 2015), our sample had residual mood symptoms limiting conclusions about the extent to which the group differences are due to trait or state. Importantly, the core finding of excessive striatal tracking of momentum-biased RPEs was independent of current mood symptoms, indicating that this may be a core feature of bipolar disorder. In contrast, only the functional connectivity findings were modulated by clinical state, perhaps indicating that the putative regulation mechanism we outlined above is further impaired by clinical state. Further research is needed to assess whether divergent functional connectivity is involved in the escalation of manic symptoms. Finally, our sample were receiving psychiatric medication. Whilst previous reviews have concluded that medications pose minimal confounds for neuroimaging studies (Hafeman et al., 2012; Phillips et al., 2008), this could nonetheless have reduced our power to detect group differences in our study. On the other hand, we did not include, by design, participants receiving antidopaminergic medications. Given the dopaminergic innervation of the loci we identified in the present study, it will be intriguing to evaluate whether the efficacy of this class of medication (Ashok et al., 2017; Kishi et al., 2021) is via ameliorating of momentum-biased RPE signaling.

In conclusion, we identify excessive striatal tracking of momentum-biased RPEs as a core mechanism in bipolar disorder, consistent with theoretical accounts (Mason et al., 2017). We additionally found divergent striatal-insular coupling during periods of upward momentum, as a putative marker for poorer contextualisation of reward perception, and exacerbation of manic symptoms. This sets the stage for evaluating this mechanism as a novel target that might be amenable to intervention in bipolar disorder.

## Funding

LM: MRC Clinician Scientist Fellowship.

HM: The author received no funding for the research, authorship and publication of this article.

## Competing interests

The authors declare that they have no competing interests.

## Notes

### Competing Interest Statement

The authors have declared no competing interest.

