## Supplementary Materials for "Misperceiving momentum: computational mechanisms of biased striatal reward prediction errors in bipolar disorder"

This file includes:

- Supplementary Formula
- Supplementary Tables
- Supplementary Results
- Supplementary Discussion

### Supplementary Formula

#### Unbiased reinforcement-learning model

In the unbiased model, reward expectations were formalised as the net expected value ($EV$) of the possible outcomes; that is the probability and value of winning the sum of money at stake combined with the probability and value of losing this stake. Reward prediction error ($RPE$), was operationalised as the difference between the actual outcome value ($r$) obtained and the expected value i.e. $RPE = r - EV$.

#### Momentum-biased reinforcement-learning model

To account for effects of momentum on each trial ($m_{(t)}$) on valuation, the unbiased model is modified to compute momentum-biased RPEs using perceived momentum-biased outcome value ($r_{biased}$) instead of the objective outcome ($r$) (Eldar & Niv, 2015).

$$r_{biased} = r * f^{m} \quad (1)$$

$$RPE_{biased} = r_{biased} - EV \quad (2)$$

Here, $f$ is the momentum bias parameter that indicates the direction and degree of momentum bias. If $f$ = 1, momentum does not bias the perception of reward or expected value. With $f$ > 1, momentum exerts positive feedback i.e. reward is perceived as larger in a good mood and smaller in a bad mood, whereas the reverse is true with 0< $f$ <1 would correspond to a negative feedback on reward value. In our present study, we set $f$ to 1.2, based on the average $f$ derived from Eldar and Niv (2015)'s sample of healthy participants who scored relatively high on the Hypomanic Personality Scale (determined by a median split) (Eckblad & Chapman, 1986), a measure of trait mood instability.

Before applying momentum to $r$, we constrained it using a sigmoid function, allowing it to take values between -1 and 1:

$$m = \tanh(m) \quad (3)$$

As per Eldar and Niv (2015)'s model, we quantified momentum on each trial, ($m_{(t)}$), as the product of momentum at the beginning of the trial, ($m_{(t-1)}$), and $RPE$ on the current trial, $RPE_{(t)}$.

$$m_{(t)} = m_{(t-1)} + RPE_{(t)} \qquad (4)$$

Unlike Eldar and Niv (2015)'s study, we did not use subjective ratings of mood to as a confirmatory check of how well the model captures participants' self-reported mood during the task. However, Eldar and Niv (2015) have confirmed the validity of this model; that it outperformed the unbiased and other reinforcement-learning models and explained participants' trial-by-trial choices and subjective mood ratings well. Hence, we assume that the model fits well here.

Given that in our task, the probability and magnitude of outcomes were fixed and made explicit in each trial, the RPEs do not have utility for updating expectations. Hence, the choice data in our task was not informative to infer each participant's learning rate, mood-update rate, which quantifies how quickly mood is updated from trial-to-trial (see below), momentum bias parameter ($f$), and momentum ($m$). Multiple studies have shown that RPEs are still tracked in tasks with minimal learning components (Rutledge et al., 2017; Rutledge et al., 2014). Given that we cannot infer mood-update rate from our task, we imposed a group-level mood-update rate on all participants, which may not fully capture inter-individual variability.

### Supplementary Tables

| Predictor | $B$ | 95% CI for $B$ | $\beta$ | $t$ | $p$ |
|---|---|---|---|---|---|
| Intercept | 8.94 | [-8.30, 26.15] | | 1.10 | .29 |
| MAS | -3.88 | [-11.12, 3.36] | -.28 | -1.13 | .27 |
| HAMD-17 | .00 | [-.78, .78] | .00 | .01 | .99 |
| BIS-11 | -.06 | [-.30, .17] | -.14 | -.58 | .57 |

**Table S1. Linear regression coefficients with manic and depressive symptoms and impulsive traits as predictors of ventral striatal activity modulated by momentum-biased RPE in participants with bipolar disorder.** $R^2$ = .12, $n$ = 21, $F(4,20)$ = .80, $p$ = .52. MAS = Bech-Rafaelsen Mania Scale; HAMD-17 = Hamilton Depression Rating Scale; BIS-11 = Barratt Impulsiveness Scale; CI = confidence interval.

| Predictor | $B$ | 95% CI for $B$ | $\beta$ | $t$ | $p$ |
|---|---|---|---|---|---|
| Intercept | -.92 | [-1.90, .04] | | -2.02 | .06 |
| MAS | -.40 | [-.80, .01] | -.45 | -2.08 | .05 |
| HAMD-17 | -.01 | [-.05, .03] | -.11 | -.51 | .62 |
| BIS-11 | .01 | [.00, .03] | .47 | 2.26 | .04 |

**Table S2. Linear regression coefficients with manic and depressive symptoms and impulsive traits as predictors of ventral striatal-left anterior insular functional connectivity modulated by momentum-biased RPE in participants with bipolar disorder.** $R^2$ = .32, $n$ = 21, $F(4,20)$ = 2.72, $p$ = .08. MAS = Bech-Rafaelsen Mania Scale; HAMD-17 = Hamilton Depression Rating Scale; BIS-11 = Barratt Impulsiveness Scale; CI = confidence interval.

### Supplementary Results

To further understand why only individuals with bipolar disorder exhibit momentum-biased striatal RPEs and whether this is driven by the mood momentum signals we identified in insula activation, we quantified the coupling between these two regions, and how it is affected by momentum of changes in reward i.e. we included model-estimated momentum values as a separate parametric regressor in functional connectivity analyses (see Supplementary Formula above for details on how model-estimated momentum values are computed).

The bipolar disorder group diverged from the healthy control group in how their striatal-left anterior insular connectivity is modulated by perception of increased momentum [$t(40) = 3.14$, $p = .0016$] with only healthy controls showing a significant modulation [Control group: $t(20) = 3.58$, $p = .00046$; Bipolar group: $t(20) = -1.04$, $p = .15$] (Figure S1).

Neither manic and depressive symptoms nor impulsive traits significantly predicted striatal-left insular functional connectivity modulated by periods of higher momentum (Table S3).

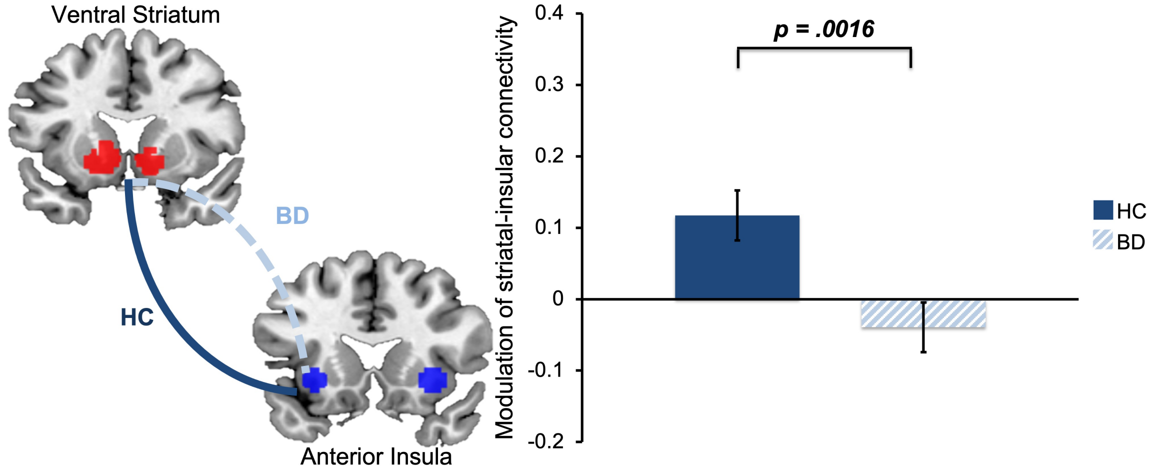


**Figure S1. Stronger modulation of ventral striatal-left anterior insular functional connectivity by periods of higher momentum in healthy controls than participants with bipolar disorder.** Bilateral ventral striatum-left anterior insular functional connectivity modulated by periods of momentum during outcome was significantly different between groups (error bars: standard error of the mean). The solid line represents significant modulation in the healthy control group (HC) and dashed line represents non-significant modulation in the bipolar disorder group (BD).

| Predictor | $B$ | 95% CI for $B$ | $\beta$ | $t$ | $p$ |
|---|---|---|---|---|---|
| Intercept | -.38 | [-.99, .24] | | -1.30 | .21 |
| MAS | -.01 | [-.27, .25] | -.02 | -.07 | .94 |
| HAMD-17 | -.01 | [-.03, .02] | -.11 | -.45 | .66 |
| BIS-11 | .01 | [-.00, .01] | .30 | 1.21 | .24 |

**Table S3. Linear regression coefficients with clinical symptoms and traits as predictors of ventral striatal-left anterior insular functional connectivity modulated by momentum-biased RPE in participants with bipolar disorder.** $R^2$ = .088, $n$ = 21, $F(4,20)$ = .55, $p$ = .66. MAS = Bech-Rafaelsen Mania Scale; HAMD-17 = Hamilton Depression Rating Scale; BIS-11 = Barratt Impulsiveness Scale; CI = confidence interval.

### Supplementary Discussion

The supplementary results reported here extend the main analyses by exploring the influence of model-estimated momentum values on ventral striatal-left anterior insular functional connectivity. These findings provide insight into how these neural processes are affected in periods of higher and lower momentum. Only healthy controls, and not participants with bipolar disorder, increased striatal-insular coupling in response to higher momentum of recent outcomes. This corroborates our main findings of stronger striatal-insular functional connectivity in healthy controls compared to participants with bipolar disorder as momentum-biased RPEs become more positive, which occurs primarily during upward momentum.

As we did not find that ventral striatal activity tracked momentum-biased RPE in healthy controls in our main findings, we reason that the significant positive modulation of striatal-insular connectivity by momentum-biased RPE could arise from changes in the momentum signal: i.e. instances where momentum-biased RPE deviates from zero correspond to instances when momentum similarly becomes strongly positive or negative. We speculate that greater ventral striatal coupling with the anterior insula helps to contextualise reward perception and reduces the chances of misperceiving the likelihood of getting future rewards from the environment. The lack of contextualisation via the striatal-insular coupling in participants with bipolar disorder could therefore result in a greater propensity to misperceive cues in the environment when mood is strongly positive or elevated, which could result in recursive cycles where expectations of reward, moods and behaviours escalate to extremes.
